# Brain-wide structural and functional disruption in mice with oligodendrocyte specific *Nf1* deletion is rescued by inhibition of NOS

**DOI:** 10.1101/2020.03.31.016089

**Authors:** Jad Asleh, Ben Shofty, Nadav Cohen, Alexandra Kavushansky, Alejandro López-Juárez, Shlomi Constantini, Nancy Ratner, Itamar Kahn

**Affiliations:** Department of Neuroscience, Rappaport Faculty of Medicine and Institute, Technion – Israel Institute of Technology, Haifa, Israel 3525422; Department of Pediatric Neurosurgery, and the Gilbert Israeli NF Center, Tel-Aviv Medical Center, and the Sackler Faculty of Medicine, Tel-Aviv University, Tel-Aviv, Israel 6423906; Division of Experimental Hematology and Cancer Biology, Cincinnati Children’s Hospital Medical Center, University of Cincinnati College of Medicine, Cincinnati, OH 45229, USA

**Author notes:** Department of Health and Biomedical Sciences, University of Texas Rio Grande Valley, TX 78520, USA. These authors contributed equally (Jad Asleh, Ben Shofty). Corresponding authors (N.R.); (I.K.).

## Abstract

Neurofibromin gene (*NF1*) mutation causes Neurofibromatosis type 1 (NF1), a disorder in which brain white matter deficits identified by neuroimaging are common, yet of unknown cellular etiology. In mice, *Nf1* loss in adult oligodendrocyte causes myelin decompaction, and increases oligodendrocyte nitric oxide (NO) levels. Nitric oxide synthase (NOS) inhibitors rescue this pathology. Whether oligodendrocyte pathology is sufficient to affect brainwide structure and account for NF1 imaging findings is unknown. Here, we show that *Nf1* gene inactivation in adult oligodendrocytes (Plp-*Nf1*^fl/+^ mice) results in a motor coordination deficit. Magnetic resonance imaging in awake mice shows that fractional anisotropy is reduced in Plp-*Nf1*^fl/+^ corpus callosum and that interhemispheric functional connectivity in motor cortex is also reduced, consistent with disrupted myelin integrity. Further, NOS-specific inhibition rescued both measures. These results demonstrate that oligodendrocyte defects account for aspects of brain dysfunction in NF1, which can be identified by neuroimaging and ameliorated by NOS inhibition.

**Significance statement:** This study assesses the effects of myelin decompaction on motor behavior and brain-wide structural and functional connectivity, and the effect of nitric oxide synthase (NOS) inhibition by N-nitro-L-arginine methyl ester (L-NAME) on these imaging measures. We report that inducible oligodendrocyte-specific inactivation of the *Nf1* gene, which causes myelin decompaction, results in reduced motor coordination. Using diffusion-based MRI we show reduced myelin integrity and using functional MRI we show reduced functional connectivity in awake passive mice. L-NAME administration results in rescue of the pathology at the mesoscopic level using imaging procedures that can be directly applied to humans to study treatment efficacy in clinical trials.

## Introduction

Neurofibromatosis type 1 (NF1) is one of the most common autosomal dominant diseases in humans, present in 1 in 2000–3000 births (Gutmann et al., 2017; Shofty et al., 2015). NF1 results from mutation in a single allele of the Neurofibromin 1 (*NF1*) gene. This can cause manifestations including skin hyperpigmentation and susceptibility to tumors of the CNS and PNS that correlate with biallelic *NF1* loss of function (Gutmann et al., 2017). Individuals with NF1 are also at risk of non-neoplastic brain disease manifestations that include attention and learning deficits, autism-like features (Baudou et al., 2020; Garg et al., 2015), and delayed acquisition of motor skills (Levine et al., 2006). NF1 children often display impairments in fine and/or gross motor function, and 40% receive remedial education to improve their cognitive, social, and motor performance (Johnson et al., 2010; Krab et al., 2008, 2011).

White matter abnormalities are frequent in NF1, and can be associated with cognitive impairment (reviewed in Baudou et al., 2020). For example, children with NF1 often show enlargement of the corpus callosum, the brain white matter tract that spans the brain hemispheres, and contains oligodendrocytes and myelinated neural axons (Cutting et al., 2002; Dubovsky et al., 2001). NF1 brain MRI findings include diffuse and focal areas of T2 hyper-intensity, speculated to represent myelin defects (Cutting et al., 2002; Pride et al., 2010; Williams et al., 2009). Diffusion tensor imaging (DTI) has been used to examine white matter microstructure in NF1. Fractional anisotropy (FA) is higher in heavily myelinated fiber tracts than in other brain regions (Feldman et al., 2010), and is reduced in NF1 children compared to typically developing children; 60–70% of NF1 children exhibit abnormalities in white matter, including reduced FA (Aydin et al., 2016; Brown et al., 2010; Karlsgodt et al., 2012; Koini et al., 2017; Nemmi et al., 2019; van der Vaart et al., 2016). All these changes lack treatment.

A histological evaluation of three NF1 brains suggested myelin edema to explain vacuolar and spongiform findings (DiPaolo et al., 1995). Consistent with the possibility that altered myelin exists in the NF1 brain, inactivation of a single *Nf1* allele in adult oligodendrocytes caused progressive myelin decompaction (López-Juárez et al., 2017; Mayes et al., 2013). Numbers of myelin lamellae were unchanged, and demyelination was not observed. Inflammation was not detected. The nitric oxide synthase (NOS) inhibitor L-arginine methyl ester (L-NAME) rescued myelin decompaction in the mouse model. In humans, whether behavioral/cognitive deficits result from *Nf1* mutation/loss in neurons, glial cells, or a combination is not defined (Gutmann et al., 2012).

Oligodendrocyte pathology is associated with decreased motor performance in humans and in mice (Kato et al., 2020; Mandolesi et al., 2019; Nomura et al., 2019). Changes in myelin itself can also correlate with altered behaviors (Franco-Pons et al., 2007; Scott et al., 2019). DTI and functional connectivity MRI (resting-state fMRI [rs-fMRI]) are used to assess structural integrity and functional connectivity in humans and rodents (Bergmann et al., 2016; Dodero et al., 2013; Grandjean et al., 2014a; Liska et al., 2015; Sforazzini et al., 2016; Zerbi et al., 2015). In NF1, functional connectivity abnormalities correlate with cognitive, social and/or behavioral deficits in human and *Nf1*^+/-^ mice (Ibrahim et al., 2017; Loitfelder et al., 2015; Shofty et al., 2019; reviewed in Baudou et al., 2020). Using human MRI techniques, with mouse-specific adaptations allowing scanning animals in an awake passive state, probing macroscopic brain-wide changes non-invasively over time bridges the microscopic and macroscopic levels and bridges basic research and clinical trials (Buckner et al., 2013; Mori and Zhang, 2006).

Here, we tested whether oligodendrocyte-specific *Nf1* mutation causes brain-wide structural and/or functional changes that impact motor areas of the brain/motor coordination, which might be rescued by NOS inhibition. Using DTI we found that Plp-*Nf1*^fl/+^ mice display reduced FA across the brain. Using rs-fMRI we identified disrupted functional connectivity in motor areas connected via the corpus callosum that were rescued by brief inhibition of NOS.

## Results

We induced oligodendrocyte-specific *Nf1* inactivation in mice and compared structural and functional differences from littermate controls (WT; *Plp-Cre* or *Nf^fl/+^*). We induced *Nf1* inactivation in eight-week-old adult mice, and 8 to 12 months later evaluated motor coordination using an accelerating rotating rod, and brain-wide connectivity using both DTI and rs-fMRI, and then evaluated the impact of inhibition, using L-NAME, on brain-related measures (**Figure 1**).

**Figure 1.**
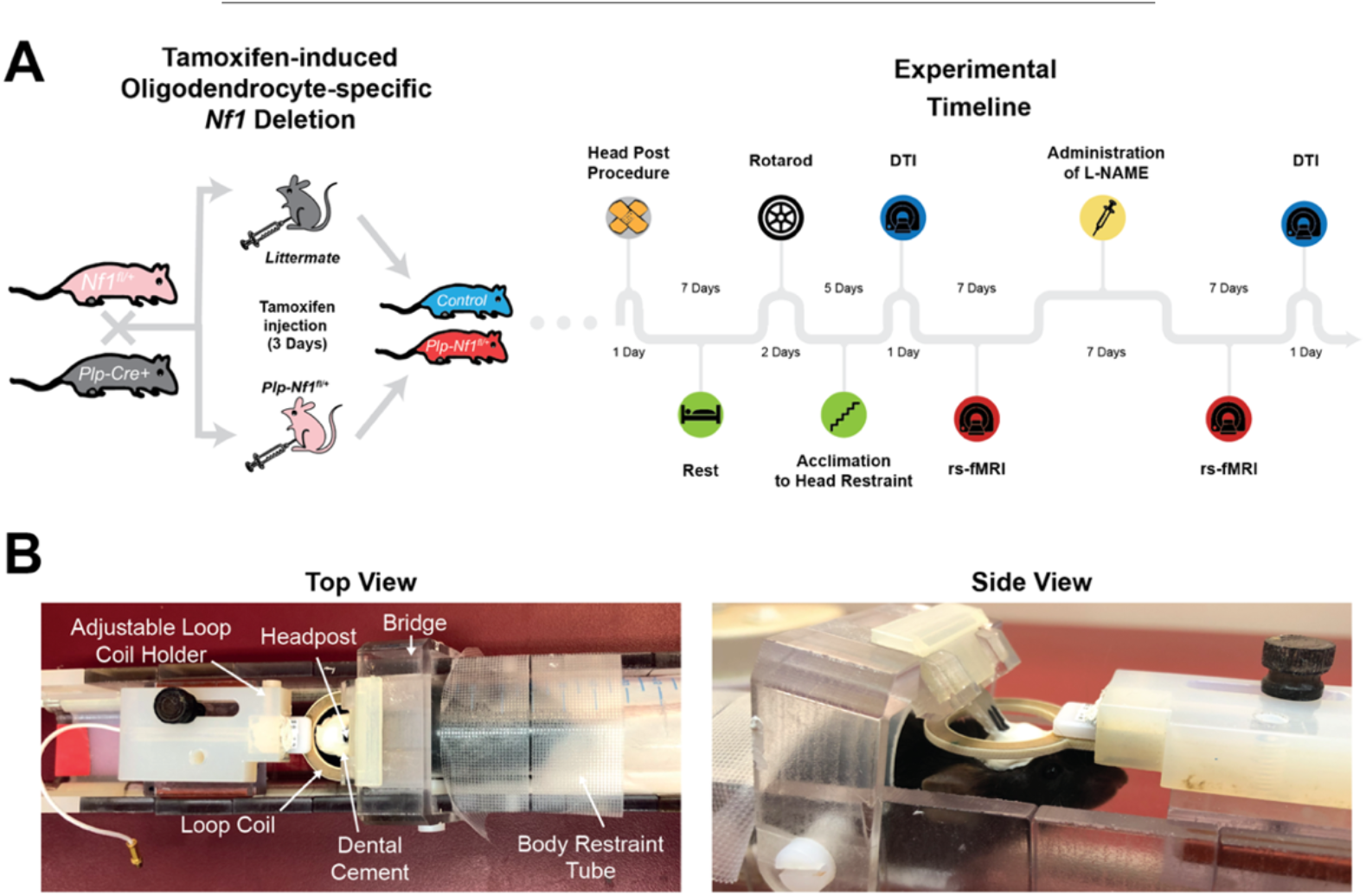
Experimental setup for whole-brain imaging of Plp-*Nf1*^fl/+^ animals. (A) Oligodendrocyte-specific *Nf1* deletion was achieved by crossing of Plp-Cre and *Nf1*^fl/+^ mice. Plp-*Nf1*^fl/+^ and littermate animals received a three-day tamoxifen treatment (twice a day) to induce the deletion in Plp-*Nf1*^fl/+^ mice and create littermate controls (Plp-*Nf1*^+/+^). The experimental timeline is shown for mice that underwent full evaluation including behavior, as well as structural and functional imaging. (B) A custom-built 3D-printed modular cradle allows head-fixation for functional scans of awake passive mice. The same cradle allows the addition of a mouse-compatible mask that can be placed around the animal’s nose and mouth to maintain constant anesthesia of the head-fixed mice for long structural scans.

To evaluate motor performance and coordination we used the accelerating rotating rod experiment over two days (four runs per day; control = 14; Plp-*Nf1*^fl/+^ = 15). Repeated measures ANOVA (**Figure 2A**) showed a main effect of genotype (*F_1,27_* = 6.3, *p* = 0.018), time (*F_1,27_* = 38.4, *p* < 0.001) and interaction (*F_1,27_* = 4.65, *p* = 0.04). Interestingly, Plp-*Nf1*^fl/+^ mice demonstrated impaired initial performance (first run; Mann-Whitney U test: *U* = 31, *p* = 0.0013) in the test and then caught up to controls (last run; *U* = 89, *p* = 0.4956). To quantify this phenomenon, we adapted a previously used analysis (Rothwell et al., 2014). Briefly, we performed a linear regression from the first run until the run where the animal first reached maximal performance (top speed of 40 revolutions per minute). From the linear regression we calculated the intercept and slope which reflect initial motor coordination (**Figure 2B**) and learning rate (**Figure 2C**), respectively. Plp-*Nf1*^fl/+^ mice showed reduced initial motor coordination (*U* = 42; *p* = 0.006), but normal learning rate (*U* = 73, *p* = 0.17). Taken together, Plp-*Nf1^fl/+^* animals show differential performance on the rotating rod with impaired initial motor coordination but normal learning rate, leading them to learn the task by the end of the eighth run and then perform at the same level as the controls.

**Figure 2.**
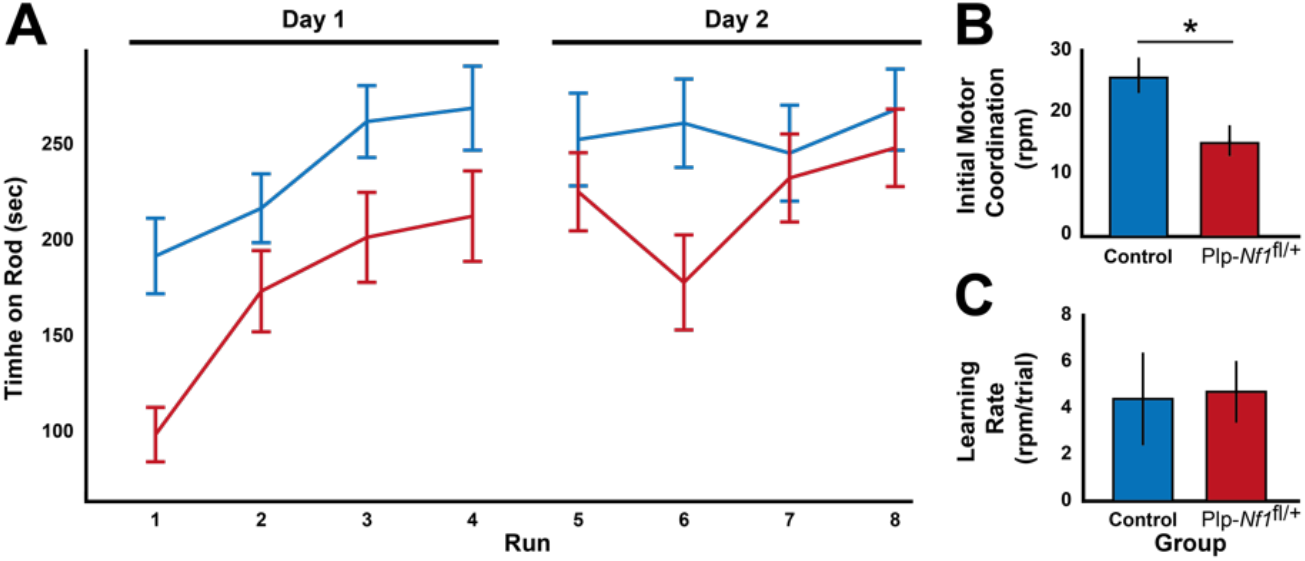
Plp-*Nf1*^fl/+^ animals show differential performance on the rotating rod with impaired initial motor coordination but normal learning rate. (A) Average time on rod is plotted for each group on consecutive testing days. Repeated measures ANOVA demonstrates an overall significant difference between Control (*n* = 14) and Plp-*Nf1*^fl/+^ (*n* = 15). (B) Initial motor coordination is plotted for Plp-*Nfl^+^* demonstrating impaired initial motor coordination (Mann-Whitney U Test: *\*p* < 0.05). (C) Learning rate is equivalent for both groups.

To determine if brain-wide genotype-specific changes in white matter integrity in this model alter fractional anisotropy (FA), we scanned controls and Plp-*Nf1*^fl/+^ mice using DTI. We calculated and co-registered FA maps of each animal onto a common atlas, and then compared the genotypes by performing a whole-brain voxel-based two-sample Student’s *t*-test. We observed a widespread, mostly symmetrical, reduction in FA across the brain (**Figure 3A**), similar to that reported in NF1 patients. We focused on selected white matter tracts as regions of interest (ROIs) to quantify effects. In order to evaluate the effects of treatment, some of the animals (control = 8; Plp-*Nf1*^fl/+^ =10) underwent intraperitoneal administration of L-NAME for seven consecutive days, followed by a repeated DTI scan to examine the same ROIs (**Figure 3B**). We observed a reliable recovery of FA to control levels in six out of seven regions including the brachium of superior colliculus (*Z* = 2.31, *p* = 0.01), fasciculus reticulata (*Z* = 2.03, *p* = 0.02), internal capsule (*Z* =1.75, *p* = 0.04), medial lemniscus (*Z* = 1.89, *p* = 0.03), posterior commissure (*Z* = 2.45, *p* = 0.007) and the anterior corpus callosum (*Z* = 2.45, *p* = 0.007), and a trend for recovery in the optic nerve (*Z* = 1.47, *p* = 0.07). When considered with a conservative correction for multiple comparisons using the Bonferroni method, the anterior corpus callosum and posterior commissure remained statistically significant at alpha level below 0.05. Taken together, the voxel-based analysis of FA shows white matter regions of marked difference between Plp-*Nf1*^fl/+^ and controls, with rescue to normal FA levels after inhibition of NOS.

**Figure 3.**
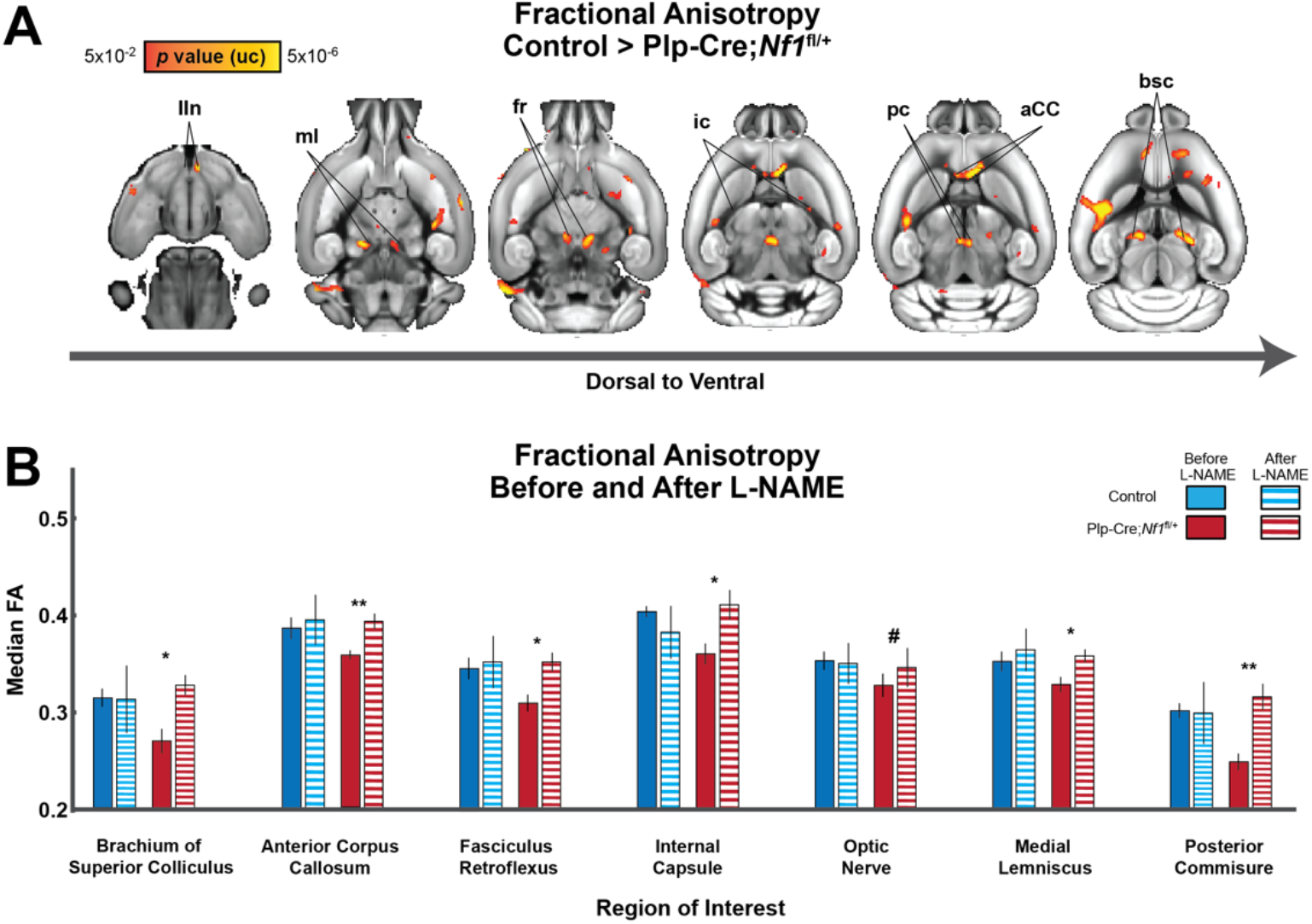
Brain-wide white matter changes in PLP mice show significant differences that are rescued by L-NAME treatment. (A) Voxel-based statistical parametric T map demonstrates reliable differences between Plp-*Nf1*^fl/+^ (*n* = 10) and Control (*n* = 11) before L-NMAE treatment in multiple white matter structures (aCC = anterior corpus callosum; bsc = brachium of superior colliculus; fr = fasciculus retroflexus; int = internal capsule; IIn = optic nerve; ml = medial lemniscus; pc = posterior commissure). (B) Response of selected brain structures to pharmacological inhibition of NOS (by L-NAME treatment) in both genotypes. (Wilcoxon signed rank test: # *p* < 0.1, * *p* < 0.05, and ** *p* < 0.01).

Based on our whole-brain DTI analysis, which showed structural damage in Plp-*Nf1*^fl/+^ mice, together with our previous evidence of disrupted white-matter integrity in this model (López-Juárez et al., 2017; Mayes et al., 2013), we hypothesized that Plp-*Nf1*^fl/+^ mice would also show disrupted functional connectivity. To test this hypothesis, we acclimated 18 animals (control = 7; Plp-*Nf1*^fl/+^ = 11; some DTI sessions were excluded due to image distortions) for awake head-fixed rs-fMRI. We scanned each mouse for 5–6 sessions (one session/day), then administered L-NAME for seven consecutive days, and then scanned for six additional sessions. We focused on the motor cortex as the anterior corpus callosum connectivity exhibited significantly reduced FA; the aCC contains axons related to the somatomotor system connecting the two hemispheres, and the integrity of this system is reflected in rotarod performance (Cao et al., 2015). As expected, the control group showed homotopicity (symmetric bilateral correlations), a hallmark of rs-fMRI in humans and rodents (Bergmann et al., 2016; Liska et al., 2015; Panzeri et al., 2017; Sforazzini et al., 2016; Shofty et al., 2019). In contrast, Plp-*Nf1*^fl/+^ mice showed reduced interhemispheric connectivity (**Figure 4A, B**; Mann-Whitney U test: *U* = 15, *p* = 0.037). This reduced connectivity was rescued by L-NAME administration (**Figure 4B**; Wilcoxon signed-rank test: *Z* = 2.134, *p* = 0.033). Collectively, these results indicate that the inhibition of NOS rescues both structural integrity and functional connectivity in Plp-*Nf1*^fl/+^ animals.

To exclude the trivial possibility that increased NO in Plp-*Nf1*^fl/+^ animals causes a global change in brain neurovascular coupling that account for these changes in the blood oxygenation level-dependent (BOLD) signal read out by fMRI, we tested whether the Plp-*Nf1*^fl/+^ group shows a non-specific difference in all interhemispheric connectivity, relative to controls. We divided the cortex into six modules based on Harris et al. (2018), further dividing the somatosensory and motor cortices such that the motor cortex is defined as MOp and somatosensory as all remaining seeds. We calculated the average interhemispheric connectivity between seeds within each module for each animal, then compared each module between genotypes (Mann-Whitney U test). No difference was observed in the anterolateral (U = 38, *p* = 0.5), somatosensory (U = 26, *p* = 0.139), visual (U = 24, *p* = 0.102), medial (U = 30, *p* = 0.234) or temporal (U = 27, *p* = 0.16) portions of the cortex. We observed a trend in prefrontal cortex (U = 20, *p* = 0.052) and a significant difference in motor cortex (U = 19, *p* = 0.032). Within the somatomotor system, rs-fMRI changes were restricted to transcallosal connections (Wilcoxon signed-rank test: *Z* = 74, *p* = 0.54). Thus, the observed changes in interhemispheric connectivity are not global and reflect the specific structural changes quantified using DTI.

Reduced FA, an indicator of myelin integrity, is predicted to affect interhemispheric cortical connectivity, which is mediated by axons crossing between sides of the brain in the corpus callosum (Roland et al., 2017; Schroeter et al., 2017). We therefore tested whether the observed FA is correlated with functional connectivity in mice that underwent both DTI and rs-fMRI; Plp-*Nf1*^fl/+^ mice but not controls showed a significant correlation between FA and rs-fMRI results (control: *r* = 0.36, *p* = 0.48; Plp-*Nf1*^fl/+^: *r* =0.9, *p* = 0.005), before L-NAME administration (**Figure 4C**; Control = 6; Plp-*Nf1*^fl/+^ = 7). It is important to exclude the possibility that the apparent rescue of reduced interhemispheric connectivity in Plp-*Nf1*^fl/+^ mice is caused by a global effect on neurovascular coupling caused by L-NAME. L-NAME blocks all NOS isoforms, thereby reducing NO and impacting neurovascular coupling, and therefore potentially the rs-fMRI signal. To exclude this possibility, we tested whether functional connectivity changed in the six cortical modules (Harris et al. 2018) in controls before and after L-NAME. We observed no difference in functional connectivity (repeated measures ANOVA; *F_1,6_* = 0.71, *p* = 0.43), suggesting that L-NAME at this dose and for this time in-and-of-itself does not introduce a global effect. Thus, Plp-*Nf1*^fl/+^ mice treated with L-NAME show specific changes in transcallosal motor system connectivity, which are detectable by rs-fMRI. These results are consistent with the demonstrated effects of L-NAME on FA in Plp-*Nf1*^fl/+^ mice shown above.

**Figure 4.**
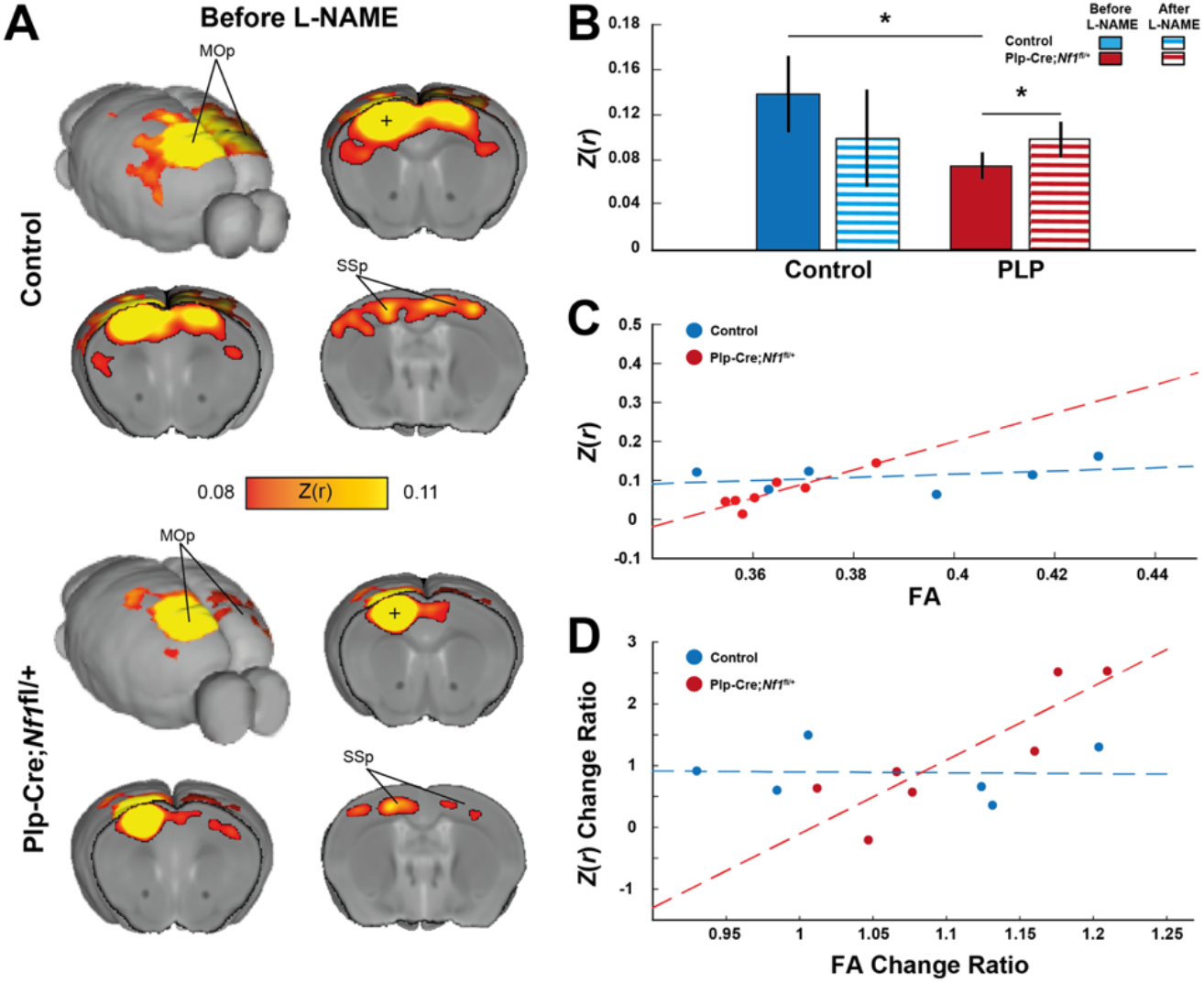
Reduced brain-wide connectivity in the motor network is rescued by L-NAME treatment. (A) Group average correlation maps demonstrate reduced interhemispheric connectivity between Control (top) and Plp-*Nf1*^fl/+^ (bottom) animals (*Z*(*r*) = Fisher’s *Z* transformed *r*; MOp = primary motor cortex; SSp = primary somatosensory cortex). (B) Correlation of seed-to-seed analysis in the primary motor cortex (denoted as a plus sign in the maps shown in A) are plotted for each group before and after L-NAME treatment, demonstrating reduced connectivity between Control and Plp-*Nf1*^fl/+^ that is rescued by inhibition of NOS. (C) FA in the corpus callosum is plotted as a function of interhemispheric functional connectivity of the motor cortex. The dashed lines represent linear fit for Control (blue) and Plp-*Nf1*^fl/+^ (red). (D) Structural and functional change due to treatment are plotted. FA change ratio defined as 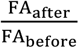 and functional connectivity *Z*(*r*) change ratio defined as 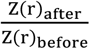 are plotted. Dashed lines represent linear fit for Control and Plp-*Nf1*^fl/+^.

Changes in FA during normal maturation, aging and disease are thought to at least partially reflect changes in myelin structure (Geeraert et al., 2019; Teipel et al., 2010). However, there was no significant correlation between FA and rs-fMRI in either group (all: *r* = 0.43, *p* = 0.14; control: *r* = 0.45, *p* = 0.39; Plp-*Nf1*^fl/+^: *r* = 0.65, *p* = 0.12). Therefore, we tested whether the changes in FA observed after L-NAME treatment result in a proportional change in functional connectivity. We observed a trend for overall correlation (**Figure 4D**; *r* = 0.51, *p* = 0.07). When considered separately, Plp-*Nf1*^fl/+^ mice showed a significant correlation (*r* = 0.87, *p* = 0.01) but controls did not (*r* = −0.0332, *p* = 0.95). We conclude that Plp-*Nf1*^fl/+^ animals with a significant difference from baseline demonstrate rescue in interhemispheric functional connectivity, which correlates with the degree of improvement observed in myelin integrity of the corpus callosum based on FA measurement.

## Discussion

NF1 is a common disease with multisystem effects that include cognitive/motor impairments and neuroimaging properties that have been proposed to result from white matter abnormalities but are of unknown etiology. Previous evidence in the *PlpCre;Nf1fl/+* mouse model linked *NF1* heterozygosity in oligodendrocytes, brain myelin producing cells, to myelin decompaction. To test if *Nf1* loss in oligodendrocytes is sufficient to disrupt structural and functional myelin integrity in living animals, we investigated the effects of oligodendrocyte-specific *Nf1* deletion using two non-invasive longitudinal brain-wide imaging systems. We found that a reduction in FA, a putative marker of myelin structure integrity, is widespread in Plp-*Nf1*^fl/+^ animals and that NOS inhibition by L-NAME rescues this defect in affected white matter tracts. Following evaluation of whole-brain structural disruption using diffusion MRI, we focused a functional connectivity analysis on the somato-motor cortex, which contains neurons whose axons travel in the damaged part of the corpus callosum identified in the DTI analysis. NOS inhibition by L-NAME rescued structural integrity and functional connectivity, and the degree of rescue was correlated in the two independent measures.

Behavioral tests in mice allow measurements of motor performance and dysfunction in a tightly controlled environment and genetic background. Here, we used the rotating rod test over five days, which tests cerebellum-independent long-term motor learning. Mice with oligodendrocyte-specific *Nf1* deletion showed reduced initial motor coordination but no change in motor-based learning. The peripheral nervous system in the Plp-*Nf1*^fl/+^ mice is not detectably abnormal (Mayes et al., 2011), suggesting that this defect is CNS based. Indicating the specificity of the *Nf1* effect in oligodendrocytes, mice overexpressing Plp in oligodendrocytes showed reduced learning, not altered co-ordination, in a forelimb movement learning task (Kato et al., 2020). *Nf1* heterozygosity in mice is associated with behavioral phenotypes including visuospatial memory and learning deficits, as in humans (Silva et al., 1997). The human impairments in fine motor coordination, manual dexterity, balance and hand/eye coordination (Krab et al., 2008; North et al., 1994) are also observed in *Nf1^+/-^* mice (Robinson et al., 2010). In contrast to Plp-*Nf1*^fl/+^ animals, which exhibited a reduction in initial motor coordination, but normal rate of motor learning, wild type and *Nf1^+/-^* mice showed similar short-term motor performance, but *Nf1^+/-^* mice showed reduced long-term motor coordination/learning not observed in the Plp-*Nf1*^fl/+^ mice (van der Vaart et al., 2011). In *Nf1^+/-^* mice, all brain cells (neurons, astrocytes, and oligodendrocytes) have reduced *Nf1*. Our finding that motor learning is intact in Plp-*Nf1*^fl/+^ mice is consistent with neuron-driven NF1-associated learning impairment, and an oligodendrocyte contribution to motor performance.

Resting-state fMRI measures slow coherent intrinsic fluctuations in oxygenated blood levels that are commonly observed between brain areas connected by monosynaptic or polysynaptic anatomic connections (Buckner et al., 2013). Damage to white-matter disrupts functional coupling between regions, suggesting that intact anatomy is necessary for functional coupling (Buckner et al., 2013; Magnuson et al., 2013). Here, we observed structural damage in the corpus callosum, the white matter tract that connects the two hemispheres, and disrupted functional connectivity in the motor areas connected via this structure in Plp-*Nf1*^fl/+^ mice. These findings are consistent with the motor co-ordination deficits in these mice being CNS, oligodendrocyte-driven.

We found that Plp-*Nf1*^fl/+^ mice display reduced FA across the brain, as measured by DTI. While the FA change is global, it was most significant in the anterior corpus callosum. Importantly, inhibition of NOS rescued FA in almost all regions examined. Thus, a readout of myelin integrity recovers in adult animals after only seven days of treatment, consistent with a similar rapid rescue of myelin integrity following NOS inhibition in this model measured by electron microscopy (López-Juárez et al., 2017). This rapid response suggests that effects of oligodendrocyte precursor cells on structural change, critical for motor learning, may not be necessary in this setting (Bells et al., 2019; Monje, 2018). Interestingly, control mice that received L-NAME, showed increased FA variability, indicating a heterogeneous response of the normal CNS to treatment, consistent with the idea that an oxidant response contributes to normal FA signal. A similar differential response of wild type mice to L-NAME was noted in motor behavior tests (Mayes et al., 2013).

We observed a correlation between the extent of change in structural and functional MRI measures, and between their absolute values. Functional connectivity and structural integrity each showed significant rescue before L-NAME, but a structure-function correlation was not preserved after L-NAME treatment. Both functional and structural imaging were acquired after treatment, but these tests were carried out sequentially; functional imaging was acquired closer to the treatment period and structural imaging acquired subsequently. These measures might become correlated after NOS inhibition by altering drug schedule, dose, or administration of different drugs. It is also possible that changes in myelination and axonal propagation integrity (which are thought to impact functional connectivity measures) are not rescued at the same rate. Additional studies are needed to characterize the timing of NOS inhibition on motor behavior and structural and functional MRI in NF1 patients to distinguish these alternatives, and to estimate the efficacy of interventions.

Previous work identified aberrant functional connectivity in NF1 children and *Nf1^+/-^* mice, with a disruption in association network organization, cortico-striatal, and cortico-cortical connectivity without reduced interhemispheric connectivity (Baudou et al., 2020; Shofty et al., 2019). This discrepancy between *Nf1* gene inactivation in all brain cells and oligodendrocyte-specific *Nf1* deletion showing reduced interhemispheric connectivity may be explained by *Nf1^+/-^* mice having mutant neurons, in addition to oligodendrocytes. *Nf1^+/-^* mice show aberrant neural activity that is mimicked by *Nf1* deletion specifically in neurons (Costa et al., 2002; Cui et al., 2008; Shilyansky et al., 2010; Tomson et al., 2015). In contrast, oligodendrocyte-specific *Nf1* inactivation impairs myelin forming cells, disrupting cellular communication, indirectly influencing neurons. Thus, it appears that when *Nf1* inactivation is present in both neurons and glia the net effect is increased cortical interhemispheric connectivity. Use of Plp-*Nf1*^fl/+^ animal model allows us to capture oligodendrocyte-specific features, partially recapitulating the human NF1 condition. Collectively, the impact of myelin decompaction on function is significant, yet not the only source contributing to the cognitive/behavioral phenotypes of this complex disorder.

Both DTI and rs-fMRI are commonly used techniques in humans. DTI may be useful as an outcome measure in future clinical trials, as FA is commonly used to identify white-matter abnormalities. However, FA is not a direct measure of myelin integrity generally or myelin decompaction specifically; other factors, such as axon composition and spacing, might affect this measure (Scholz et al., 2009). With these reservations, given that myelin decompaction is rescued by administration of L-NAME in Plp-*Nf1*^fl/+^ animals on the microscopic level (López-Juárez et al., 2017; Mayes et al., 2013), we infer that FA, at least partially, reflects the integrity of myelin in this model.

Rs-fMRI networks have been observed repeatedly and reliably under the presence of anesthetic vasodilators (Grandjean et al., 2014b; Zerbi et al., 2015); anesthetic vasodilators impact signal-to-noise but network organization is preserved. Thus, we would like to propose that both FA and rs-fMRI can be used as outcome measures in clinical trials.

We note that NOS inhibition via L-NAME is not commonly used in humans. Although L-NAME administration to patients in shock did not demonstrate any significant adverse events, it causes systemic and pulmonary hypertension, and is teratogenic during pregnancy (Kiehl et al., 1997). Possible consequences of chronic L-NAME use in children are unknown. Other safer and/or specific NOS inhibitors or anti-oxidants (e.g. N-acetyl-cysteine) might be tested in pre-clinical or clinical trials, and be better tolerated (Wong and Lerner, 2015) than Ras pathway specific inhibitors (e.g., MEK inhibitors; Broman et al., 2019), and any cognitive/motor rescue anti-oxidant affords may be of value to NF1 patients.

In summary, NOS inhibition recues structural integrity and functional connectivity across the entire brain in the Plp-*Nf1*^fl/+^ animal model. Rescued myelin integrity was correlated with rescue in functional connectivity, albeit to varying degrees, and the extent of rescue in FA is a predictor of the amount of recovery observed in interhemispheric connectivity. Given that this measure of structural MRI is widely used in humans, we suggest that it may be used as an outcome measure of future clinical trials aimed at rescuing myelin decompaction.

## Materials and Methods

### Ethics statement

All animal procedures were conducted with the approval of the Institutional Animal Care and Use Committee and are in accord with the NIH Guide for the Care and Use of Laboratory Animals.

### Animals

Inducible *Plp-Cre+* (Doerflinger et al., 2003) and *Nf1*^fl/+^ (Zhu et al., 2001). Mice were maintained on a pure C57Bl/6 background, and genotyped by PCR according to publicly available protocols (Mayes et al., 2013). Tamoxifen was administered to the animals at the age of eight weeks (see below). Twenty-nine animals participated in Rotarod evaluation (Control = 14; Plp-*Nf1*^fl/+^ = 15), 19 animals in DTI (Control = 9; Plp-*Nf1*^fl/+^ = 10), and 18 animals in rs-fMRI (Control = 7; Plp-*Nf1*^fl/+^ = 11). Of the animals that participated in rs-fMRI, 13 animals participated in DTI (Control = 6; Plp-*Nf1*^fl/+^ = 7), and 14 in Rotarod (Control = 6; Plp-*Nf1*^fl/+^ = 8).

### Tamoxifen injection

The injection protocol followed previous publications (López-Juárez et al., 2017; Mayes et al., 2013). We dissolved tamoxifen (100 mg) with 1 ml of ethanol and 9 ml of sunflower seed oil (Sigma-Aldrich). At the age of eight weeks, both controls and Plp-*Nf1*^fl/+^ mice were injected with tamoxifen (100 μl) intraperitoneally, twice a day for three days, at the same time-of-day.

### N-nitro-L-arginine methyl ester (L-NAME) injection

Fresh solutions of L-NAME were prepared daily in accordance with previously published protocols (López-Juárez et al., 2017; L-NG-nitroarginine methyl ester, 100 mM in 13 PBS; Sigma-Aldrich) and injected to animals intraperitoneally (0.4 mg/kg) daily, for seven days.

### Animal surgery

Prior to any experiment, to use our in-house designed cradle, we implanted a headpost on the animal’s skull as previously described (Bergmann et al., 2016). Briefly, the animals were anesthetized with isoflurane mixed with oxygen and kept at a level of 2–3% for the entire surgery. The skin was removed and a 3D-printed plastic headpost was positioned on the animal’s skull and fixed by applying dental cement (C&B Metabond, Parkell). To prevent image distortions in MRI acquisition resulting from uneven surfaces, the cement was smoothed using a self-cured resin (General Purpose Acrylic Resin, Unifast Trad). The animals were then left to recover and injected once with antibiotics (Cefalexin 0.18 mg/10g; Norbrook) along with systemic and local analgesics (Buprenorphine 0.9 μg/10g; Bupavacaine 0.075mg/10g, respectively). Each animal was carefully monitored for the next week before entering the experiment.

### Rotating rod performance test

Gross motor coordination function was evaluated using Rotarod (Med Associates, Vermont, USA). Twenty-nine animals (Control = 14; Plp-*Nf1*^fl/+^ = 15) underwent one training and one test session, each consisting of four runs. Every run lasted for six minutes, in which the rod accelerated gradually from 4 to 40 revolutions per minute in the first five minutes and maintained at the final speed for the last minute. Time-to-fall and peak speed were recorded.

### MRI acquisition

Imaging was performed using a 9.4 tesla BioSpec 94/20 USR MRI (Bruker BioSpin MRI). A 20 mm receive-only coil (Bruker BioSpin MRI) was fixed to the cradle over the animal’s head, and the entire apparatus was inserted into the MRI bore containing an 86 mm quadrature transmit-only coil (Bruker BioSpin MRI).

### Diffusion tensor imaging (DTI) acquisition

To characterize structural connectivity, we scanned 19 animals using DTI (control = 9; Plp-*Nf1*^fl/+^ = 10); one control was excluded due to extensive image distortions. Animals were first lightly sedated and fixed to the resting-state cradle with a custom-made mask placed over their nose which allowed to maintain anesthesia throughout the DTI scanning session while their respiratory rate was monitored. Data acquisition was performed using a diffusion-weighted spin-echo echo-planar imaging (EPI) pulse sequence (TR = 7000 ms, TE = 21.68ms). We used 30 non-colinear diffusion directions, a b_factor_ of 1000 s/mm^3^ with 11 ms duration of the diffusion gradient (δ) and a separation time (Δ) of 2.6 ms. The scan consisted of 24 slices of 400 μm thickness, with an acquisition matrix of 100 × 100 voxels and a field of view of 16 × 16 cm^2^ resulting in a resolution of 0.1 × 0.1 × 0.4 mm^3^.

### DTI preprocessing and analysis

Diffusion weighted images were preprocessed using ExplorDTI (Leemans et al., 2009) with tensors calculated using a robust estimation algorithm (Krestin et al., 2014), together with motion and eddy current correction. FA and B_0_ maps were exported to NIfTI format. Next, brain extraction was performed and a template B_0_ for both controls and Plp-*Nf1*^fl/+^ animals was created using ANTs (Avants et al., 2008). The controls’ template was registered to the Allen Brain

Atlas (Oh et al., 2014) while the Plp-*Nf1*^fl/+^ was linearly co-registered to the controls. A final diffeomorphic field from native space to the Allen Brain space for each animal was calculated and applied in one step to native scans to prevent data degradation.

Normalized FA maps were submitted to a two-sample Student’s *t*-test between controls and Plp-*Nf1*^fl/+^ animals. For ROI analysis we defined the regions manually based on the *t*-test results comparing genotypes before L-NAME, and calculated the mean voxel intensity within each region to assess rescue after treatment (Jenkinson et al., 2012).

### Resting-state fMRI (rs-fMRI) acquisition

Eighteen animals were included in the awake resting-state fMRI experiment (Control = 7; Plp-*Nf1*^fl/+^ = 11) which followed a previously published protocol (Bergmann et al., 2016). To reduce stress, the animals were first lightly sedated and fixed to the resting-state cradle. During the five-day acclimatization period, each animal was placed in the MRI and exposed to the different scanning protocols used, while the head restraint period was increased gradually from 2 to 50 minutes by the last acclimatization session. Afterwards, each animal was scanned before and after the administration of L-NAME for 6–7 daily sessions of rs-fMRI at each time period. Each scanning session included structural and functional scanning. The structural scan was acquired using a T_1_-weigthed rapid acquisition process with relaxation enhancement (RARE) sequence in coronal orientation (30 coronal slices, TE = 8.5 ms, RARE factor = 4, flip angle = 180°, in-plane resolution 150 × 150 μm^2^, slice thickness = 450 μm). Whole-brain functional scans consisted of 800 repetitions divided over four runs (200 repetitions per run). Blood oxygenation level-dependent (BOLD) imaging was achieved using spin echo-echo planar imaging (SE-EPI) sequence (TR = 2.5 s, TE = 18.398 ms, flip angle = 90°, 30 coronal slices, in-plane resolution = 150 × 150 μm^2^, slice thickness = 450 μm; the imaged volume was framed with four saturation slices to avoid wraparound artifacts).

### Rs-fMRI preprocessing and analysis

The preprocessing steps were previously reported (Bergmann et al., 2016; Kahn et al., 2011). Briefly, the first two frames were skipped to eliminate inhomogeneity artifacts and slice time corrected. To perform motion correction, the whole time-series was corrected to the average of the first 200 repetitions using rigid-body correction (Jenkinson et al., 2002) by calculating a coregistration matrix for each volume. The same averaged volume was then warped automatically, using symmetric diffeomorphic image normalization (SyN) by Advanced Normalization Tools (ANTs; Avants, Tustison, & Song, 2009), to a predefined EPI template that was preregistered to the Allen Brain Atlas. To prevent degradation of image quality, the final displacement field for each volume, including its motion correction displacement matrix, was calculated and applied on each time-corrected volume. Whole-brain and ventricle masks were calculated automatically. Data-scrubbing procedures were similar to previous literature (Bergmann et al., 2016; Power et al., 2014) with similar exclusion criteria (frame displacement threshold of 50 μm, temporal derivative mean square root area [DVARS] threshold of 150% inter-quartile range [IQR] above the 75^th^ percentile with exclusion of one frame after the detected motion). Runs with fewer than 50 frames and sessions with fewer than 300 frames were excluded; 29 out of 208 sessions were excluded due to excessive movement or image distortion (control = 11; Plp-*Nf1*^fl/+^ = 18). After data-scrubbing, the time series underwent demeaning and de-trending to remove global mean and non-intrinsic trends, together with regressing out mean ventricles and vascular signals, and six volume-specific motion parameters and their derivatives were regressed out of the signal to remove sources of spurious or regionally nonspecific variance (Grandjean et al., 2020). Finally, band-pass filtering (0.009–0.08 Hz) and smoothing using convolution with a Gaussian function (FWHM of 0.6 mm) were applied.

To estimate functional connectivity maps, we placed a region of interest with a 100 μm radius and extracted the time course from each seed. A seed to whole brain correlation map was calculated over each session of the animal and averaged to produce a mean Fisher’s *Z* transform, *Z*(*r*), map for that animal. Finally, all the animal-based maps were averaged to produce one map for each genotype that were displayed on a template registered to the Allen Brain Atlas (Dorr et al., 2008). Seed-to-seed analysis was performed by calculating Pearson’s correlation coefficient between two seeds and transforming it using Fisher’s *Z* transform to allow for cross-subject comparison.

## Acknowledgments

This work was supported by the Israel Science Foundation (770/17), and the Adelis Foundation (to I.K.), and the National Institutes of Health (1R01NS091037; to N.R. and I.K.). We thank the Technion’s Biological Core Facilities and Edith Suss-Toby for her assistance with MRI, and the Technion Preclinical Research Authority for assistance with animal care.

## References

Avants, B.B., Epstein, C.L., Grossman, M., and Gee, J.C. (2008). Symmetric diffeomorphic image registration with cross-correlation: Evaluating automated labeling of elderly and neurodegenerative brain. Med. Image Anal. 12, 26–41.

Avants, B.B., Tustison, N., and Song, G. (2009). Advanced Normalization Tools (ANTS). Insight J. 1–35.

Aydin, S., Kurtcan, S., Alkan, A., Guler, S., Filiz, M., Yilmaz, T.F., Sahin, T.U., and Aralasmak, A. (2016). Relationship between the corpus callosum and neurocognitive disabilities in children with NF-1: diffusion tensor imaging features. Clin. Imaging 40, 1092–1095.

Baudou, E., Nemmi, F., Biotteau, M., Maziero, S., Peran, P., and Chaix, Y. (2020). Can the Cognitive Phenotype in Neurofibromatosis Type 1 (NF1) Be Explained by Neuroimaging? A Review. Front. Neurol. 10, 1–11.

Bells, S., Lefebvre, J., Longoni, G., Narayanan, S., Arnold, D.L., Yeh, E.A., and Mabbott, D.J. (2019). White matter plasticity and maturation in human cognition. Glia 67, 2020–2037.

Bergmann, E., Zur, G., Bershadsky, G., and Kahn, I. (2016). The organization of mouse and human cortico-hippocampal networks estimated by intrinsic functional connectivity. Cereb. Cortex 26, 4497–4512.

Broman, K.K., Dossett, L.A., Sun, J., Eroglu, Z., and Zager, J.S. (2019). Update on BRAF and MEK inhibition for treatment of melanoma in metastatic, unresectable, and adjuvant settings. Expert Opin. Drug Saf. 18, 381–392.

Brown, J.A., Emnett, R.J., White, C.R., Yuede, C.M., Conyers, S.B., O’Malley, K.L., Wozniak, D.F., and Gutmann, D.H. (2010). Reduced striatal dopamine underlies the attention system dysfunction in neurofibromatosis-1 mutant mice. Hum. Mol. Genet. 19, 4515–4528.

Buckner, R.L., Krienen, F.M., and Yeo, B.T.T. (2013). Opportunities and limitations of intrinsic functional connectivity MRI. Nat. Neurosci. 16, 832–837.

Cao, V.Y., Ye, Y., Mastwal, S., Ren, M., Coon, M., Liu, Q., Costa, R.M., and Wang, K.H. (2015). Motor Learning Consolidates Arc-Expressing Neuronal Ensembles in Secondary Motor Cortex. Neuron 86, 1385–1392.

Costa, R.M., Federov, N.B., Kogan, J.H., Murphy, G.G., Stern, J., Ohno, M., Kucherlapati, R., Jacks, T., and Silva, A.J. (2002). Mechanism for the learning deficits in a mouse model of neurofibromatosis type 1. Nature 415, 526–530.

Cui, Y., Costa, R.M., Murphy, G.G., Elgersma, Y., Zhu, Y., Gutmann, D.H., Parada, L.F., Mody, I., and Silva, A.J. (2008). Neurofibromin regulation of ERK signaling modulates GABA release and learning. Cell 135, 549–560.

Cutting, L.E., Cooper, K.L., Koth, C.W., Mostofsky, S.H., Kates, W.R., Denckla, M.B., and Kaufmann, W.E. (2002). Megalencephaly in NF1: Predominantly white matter contribution and mitigation by ADHD. Neurology 59, 1388–1394.

DiPaolo, D.P., Zimmerman, R.A., Rorke, L.B., Zackai, E.H., Bilaniuk, L.T., and Yachnis, A.T. (1995). Neurofibromatosis type 1: Pathologic substrate of high-signal-intensity foci in the brain. Radiology 195, 721–724.

Dodero, L., Damiano, M., Galbusera, A., Bifone, A., Tsaftsaris, S.A., Scattoni, M.L., and Gozzi, A. (2013). Neuroimaging Evidence of Major Morpho-Anatomical and Functional Abnormalities in the BTBR T+TF/J Mouse Model of Autism. PLoS One.

Doerflinger, N.H., Macklin, W.B., and Popko, B. (2003). Inducible site-specific recombination in myelinating cells. Genesis 35, 63–72.

Dorr, A.E., Lerch, J.P., Spring, S., Kabani, N., and Henkelman, R.M. (2008). High resolution three-dimensional brain atlas using an average magnetic resonance image of 40 adult C57Bl/6J mice. Neuroimage 42, 60–69.

Dubovsky, E.C., Booth, T.N., Vezina, G., Samango-Sprouse, C.A., Palmer, K.M., and Brasseux, C.O. (2001). MR imaging of the corpus callosum in pediatric patients with neurofibromatosis type 1. AJNR. Am. J. Neuroradiol. 22, 190–195.

Feldman, H.M., Yeatman, J.D., Lee, E.S., Barde, L.H.F., and Gaman-Bean, S. (2010). Diffusion Tensor Imaging: A Review for Pediatric Researchers and Clinicians. J. Dev. Behav. Pediatr. 31, 346–356.

Franco-Pons, N., Torrente, M., Colomina, M.T., and Vilella, E. (2007). Behavioral deficits in the cuprizone-induced murine model of demyelination/remyelination. Toxicol. Lett. 169, 205–213.

Garg, S., Plasschaert, E., Descheemaeker, M.J., Huson, S., Borghgraef, M., Vogels, A., Evans, D.G., Legius, E., and Green, J. (2015). Autism Spectrum Disorder Profile in Neurofibromatosis Type I. J. Autism Dev. Disord. 45, 1649–1657.

Geeraert, B.L., Lebel, R.M., and Lebel, C. (2019). A multiparametric analysis of white matter maturation during late childhood and adolescence. Hum. Brain Mapp. 40, 4345–4356.

Grandjean, J., Schroeter, A., He, P., Tanadini, M., Keist, R., Krstic, D., Konietzko, U., Klohs, J., Nitsch, R.M., and Rudin, M. (2014a). Early alterations in functional connectivity and white matter structure in a transgenic mouse model of cerebral amyloidosis. J. Neurosci. 34, 13780–13789.

Grandjean, J., Schroeter, A., Batata, I., and Rudin, M. (2014b). Optimization of anesthesia protocol for resting-state fMRI in mice based on differential effects of anesthetics on functional connectivity patterns. Neuroimage 102, 838–847.

Grandjean, J., Canella, C., Anckaerts, C., Ayrancı, G., Bougacha, S., Bienert, T., Buehlmann, D., Coletta, L., Gallino, D., Gass, N., et al. (2020). Common functional networks in the mouse brain revealed by multi-centre resting-state fMRI analysis. Neuroimage 205.

Gutmann, D.H., Parada, L.F., Silva, A.J., and Ratner, N. (2012). Neurofibromatosis type 1: Modeling CNS dysfunction. J. Neurosci. 32, 14087–14093.

Gutmann, D.H., Ferner, R.E., Listernick, R.H., Korf, B.R., Wolters, P.L., and Johnson, K.J. (2017). Neurofibromatosis type 1. Nat. Rev. Dis. Prim. 3, 17004.

Harris, J.A., Mihalas, S., Hirokawa, K.E., Whitesell, J.D., Knox, J., Bernard, A., Bohn, P., Caldejon, S., Casal, L., Cho, A., et al. (2018). The organization of intracortical connections by layer and cell class in the mouse brain. BioRxiv 292961.

Ibrahim, A.F.A., Montojo, C.A., Haut, K.M., Karlsgodt, K.H., Hansen, L., Congdon, E., Rosser, T., Bilder, R.M., Silva, A.J., and Bearden, C.E. (2017). Spatial working memory in neurofibromatosis 1: Altered neural activity and functional connectivity. NeuroImage Clin. 15, 801–811.

Jenkinson, M., Bannister, P., Brady, M., and Smith, S. (2002). Improved optimization for the robust and accurate linear registration and motion correction of brain images. Neuroimage 17, 825–841.

Jenkinson, M., Beckmann, C.F., Behrens, T.E.J., Woolrich, M.W., and Smith, S.M. (2012). FSL. Neuroimage 62, 782–790.

Johnson, B.A., MacWilliams, B.A., Carey, J.C., Viskochil, D.H., D’Astous, J.L., and Stevenson, D.A. (2010). Motor proficiency in children with neurofibromatosis type 1. Pediatr. Phys. Ther. 22, 344–348.

Kahn, I., Desai, M., Knoblich, U., Bernstein, J., Henninger, M., Graybiel, A.M., Boyden, E.S., Buckner, R.L., and Moore, C.I. (2011). Characterization of the Functional MRI Response Temporal Linearity via Optical Control of Neocortical Pyramidal Neurons. J. Neurosci. 31, 15086–15091.

Karlsgodt, K.H., Rosser, T., Lutkenhoff, E.S., Cannon, T.D., Silva, A., and Bearden, C.E. (2012). Alterations in White Matter Microstructure in Neurofibromatosis-1. PLoS One 7, e47854.

Kato, D., Wake, H., Lee, P.R., Tachibana, Y., Ono, R., Sugio, S., Tsuji, Y., Tanaka, Y.H., Tanaka, Y.R., Masamizu, Y., et al. (2020). Motor learning requires myelination to reduce asynchrony and spontaneity in neural activity. Glia 68, 193–210.

Kiehl, M.G., Ostermann, H., Meyer, J., and Kienast, J. (1997). Nitric oxide synthase inhibition by L-NAME in leukocytopenic patients with severe septic shock. Intensive Care Med. 23, 561–566.

Koini, M., Rombouts, S.A.R.B., Veer, I.M., Van Buchem, M.A., and Huijbregts, S.C.J. (2017). White matter microstructure of patients with neurofibromatosis type 1 and its relation to inhibitory control. Brain Imaging Behav. 11, 1731–1740.

Krab, L.C., Aarsen, F.K., de Goede-Bolder, A., Catsman-Berrevoets, C.E., Arts, W.F., Moll, H.A., and Elgersma, Y. (2008). Impact of neurofibromatosis type 1 on school performance. J. Child Neurol. 23, 1002–1010.

Krab, L.C., De Goede-Bolder, A., Aarsen, F.K., Moll, H.A., De Zeeuw, C.I., Elgersma, Y., and Van Der Geest, J.N. (2011). Motor learning in children with neurofibromatosis type I. Cerebellum 10, 14–21.

Krestin, G.P., Heemskerk, A.M., Dudink, J., Leemans, A., Govaert, P., Reiss, I.K.M., Lequin, M.H., Pieterman, K., and Plaisier, A. (2014). Choice of Diffusion Tensor Estimation Approach Affects Fiber Tractography of the Fornix in Preterm Brain. Am. J. Neuroradiol. 35, 1219–1225.

Leemans, A., Jeurissen, B., Sijbers, J., and Jones, D. (2009). ExploreDTI: a graphical toolbox for processing, analyzing, and visualizing diffusion MR data. Proc. 17th Sci. Meet. Int. Soc. Magn. Reson. Med. 17, 3537.

Levine, T.M., Materek, A., Abel, J., O’Donnell, M., and Cutting, L.E. (2006). Cognitive Profile of Neurofibromatosis Type 1. Semin. Pediatr. Neurol. 13, 8–20.

Liska, A., Galbusera, A., Schwarz, A.J., and Gozzi, A. (2015). Functional connectivity hubs of the mouse brain. Neuroimage 115, 281–291.

Loitfelder, M., Huijbregts, S.C.J., Veer, I.M., Swaab, H.S., Van Buchem, M.A., Schmidt, R., and Rombouts, S.A. (2015). Functional connectivity changes and executive and social problems in neurofibromatosis type i. Brain Connect. 5, 312–320.

López-Juárez, A., Titus, H.E., Silbak, S.H., Pressler, J.W., Rizvi, T.A., Bogard, M., Bennett, M.R., Ciraolo, G., Williams, M.T., Vorhees, C. V., et al. (2017). Oligodendrocyte Nf1 Controls Aberrant Notch Activation and Regulates Myelin Structure and Behavior. Cell Rep. 19, 545–557.

Magnuson, M.E., Thompson, G.J., Pan, W.-J., and Keilholz, S.D. (2013). Effects of severing the corpus callosum on electrical and BOLD functional connectivity and spontaneous dynamic activity in the rat brain. Brain Connect. 4, 131011122122002.

Mandolesi, G., Bullitta, S., Fresegna, D., De Vito, F., Rizzo, F.R., Musella, A., Guadalupi, L., Vanni, V., Stampanoni Bassi, M., Buttari, F., et al. (2019). Voluntary running wheel attenuates motor deterioration and brain damage in cuprizone-induced demyelination. Neurobiol. Dis. 129, 102–117.

Mayes, D.A., Rizvi, T.A., Cancelas, J.A., Kolasinski, N.T., Ciraolo, G.M., Stemmer-Rachamimov, A.O., and Ratner, N. (2011). Perinatal or adult Nf1 inactivation using tamoxifen-inducible PlpCre each cause neurofibroma formation. Cancer Res. 71, 4675–4685.

Mayes, D.A., Rizvi, T.A., Titus-Mitchell, H., Oberst, R., Ciraolo, G.M., Vorhees, C. V., Robinson, A.P., Miller, S.D., Cancelas, J.A., Stemmer-Rachamimov, A.O., et al. (2013). Nf1 Loss and Ras Hyperactivation in Oligodendrocytes Induce NOS-Driven Defects in Myelin and Vasculature. Cell Rep. 4, 1197–1212.

Monje, M. (2018). Myelin Plasticity and Nervous System Function. Annu. Rev. Neurosci. 41, 61–76.

Mori, S., and Zhang, J. (2006). Principles of Diffusion Tensor Imaging and Its Applications to Basic Neuroscience Research. Neuron 51, 527–539.

Nemmi, F., Cignetti, F., Assaiante, C., Maziero, S., Audic, F., Péran, P., and Chaix, Y. (2019). Discriminating between neurofibromatosis-1 and typically developing children by means of multimodal MRI and multivariate analyses. Hum. Brain Mapp. 40, 3508–3521.

Nomura, T., Bando, Y., Nakazawa, H., Kanemoto, S., and Yoshida, S. (2019). Pathological changes in mice with long term cuprizone administration. Neurochem. Int. 126, 229–238.

North, K., Joy, P., Yuille, D., Cocks, N., Mobbs, E., Hutchins, P., McHugh, K., and de Silva, M. (1994). Specific learning disability in children with neurofibromatosis type 1: Significance of MRI abnormalities. Neurology 44, 878–878.

Oh, S.W., Harris, J.A., Ng, L., Winslow, B., Cain, N., Mihalas, S., Wang, Q., Lau, C., Kuan, L., Henry, A.M., et al. (2014). A mesoscale connectome of the mouse brain. Nature 508, 207–214.

Panzeri, S., Gomolka, R., Pasqualetti, M., Gozzi, A., Galbusera, A., Scattoni, M.L., Bertero, A., Barsotti, N., Sabbioni, M., and Liska, A. (2017). Homozygous Loss of Autism-Risk Gene CNTNAP2 Results in Reduced Local and Long-Range Prefrontal Functional Connectivity. Cereb. Cortex.

Power, J.D., Mitra, A., Laumann, T.O., Snyder, A.Z., Schlaggar, B.L., and Petersen, S.E. (2014). Methods to detect, characterize, and remove motion artifact in resting state fMRI. Neuroimage 84, 320–341.

Pride, N., Payne, J.M., Webster, R., Shores, E.A., Rae, C., and North, K.N. (2010). Corpus callosum morphology and its relationship to cognitive function in neurofibromatosis type 1. J. Child Neurol. 25, 834–841.

Robinson, A., Kloog, Y., Stein, R., and Assaf, Y. (2010). Motor deficits and neurofibromatosis type 1 (NF1)-associated MRI impairments in a mouse model of NF1. NMR Biomed. 23, 1173–1180.

Roland, J.L., Snyder, A.Z., Hacker, C.D., Mitra, A., Shimony, J.S., Limbrick, D.D., Raichle, M.E., Smyth, M.D., and Leuthardt, E.C. (2017). On the role of the corpus callosum in interhemispheric functional connectivity in humans. Proc. Natl. Acad. Sci. 114, 13278–13283.

Rothwell, P.E., Fuccillo, M. V., Maxeiner, S., Hayton, S.J., Gokce, O., Lim, B.K., Fowler, S.C., Malenka, R.C., and Südhof, T.C. (2014). Autism-associated neuroligin-3 mutations commonly impair striatal circuits to boost repetitive behaviors. Cell 158, 198–212.

Scholz, J., Tomassini, V., and Johansen-Berg, H. (2009). Chapter 11 - Individual Differences in White Matter Microstructure in the Healthy Brain. In Diffusion MRI, H. Johansen-Berg, and T.E.J. Behrens, eds. (San Diego: Academic Press), pp. 237–249.

Schroeter, A., Grandjean, J., Schlegel, F., Saab, B.J., and Rudin, M. (2017). Contributions of structural connectivity and cerebrovascular parameters to functional magnetic resonance imaging signals in mice at rest and during sensory paw stimulation. J. Cereb. Blood Flow Metab. 37, 2368–2382.

Scott, R., Sánchez-Aguilera, A., Van Elst, K., Lim, L., Dehorter, N., Bae, S.E., Bartolini, G., Peles, E., Kas, M.J.H., Bruining, H., et al. (2019). Loss of Cntnap2 Causes Axonal Excitability Deficits, Developmental Delay in Cortical Myelination, and Abnormal Stereotyped Motor Behavior. Cereb. Cortex 29, 586–597.

Sforazzini, F., Bertero, A., Dodero, L., David, G., Galbusera, A., Scattoni, M.L., Pasqualetti, M., and Gozzi, A. (2016). Altered functional connectivity networks in acallosal and socially impaired BTBR mice. Brain Struct. Funct. 221, 941–954.

Shilyansky, C., Karlsgodt, K.H., Cummings, D.M., Sidiropoulou, K., Hardt, M., James, A.S., Ehninger, D., Bearden, C.E., Poirazi, P., Jentsch, J.D., et al. (2010). Neurofibromin regulates corticostriatal inhibitory networks during working memory performance. Proc. Natl. Acad. Sci. 107, 13141–13146.

Shofty, B., Constantini, S., and Ben-Shachar, S. (2015). Advances in Molecular Diagnosis of Neurofibromatosis Type 1. Semin. Pediatr. Neurol. 22, 234–239.

Shofty, B., Bergmann, E., Zur, G., Asleh, J., Bosak, N., Kavushansky, A., Castellanos, F.X., Ben-Sira, L., Packer, R.J., Vezina, G.L., et al. (2019). Autism-associated Nf1 deficiency disrupts corticocortical and corticostriatal functional connectivity in human and mouse. Neurobiol. Dis. 130, 104479.

Silva, A.J., Frankland, P.W., Marowitz, Z., Friedman, E., Laszlo, G.S., Cioffi, D., Jacks, T., Bourtchuladze, R., Lazlo, G., Cioffi, D., et al. (1997). A mouse model for the learning and memory deficits associated with neurofibromatosis type I. Nat. Genet. 15, 281–284.

Teipel, S.J., Meindl, T., Wagner, M., Stieltjes, B., Reuter, S., Hauenstein, K.-H., Filippi, M., Ernemann, U., Reiser, M.F., and Hampel, H. (2010). Longitudinal Changes in Fiber Tract Integrity in Healthy Aging and Mild Cognitive Impairment: A DTI Follow-Up Study. J. Alzheimer’s Dis. 22, 507–522.

Tomson, S.N., Schreiner, M.J., Narayan, M., Rosser, T., Enrique, N., Silva, A.J., Allen, G.I., Bookheimer, S.Y., and Bearden, C.E. (2015). Resting state functional MRI reveals abnormal network connectivity in neurofibromatosis 1. Hum. Brain Mapp. 36.

van der Vaart, T., van Woerden, G.M., Elgersma, Y., de Zeeuw, C.I., and Schonewille, M. (2011). Motor deficits in neurofibromatosis type 1 mice: The role of the cerebellum. Genes, Brain Behav. 10, 404–409.

van der Vaart, T., Rietman, A.B., Plasschaert, E., Legius, E., Elgersma, Y., and Moll, H.A. (2016). Behavioral and cognitive outcomes for clinical trials in children with neurofibromatosis type 1. Neurology 86, 154–160.

Williams, V.C., Lucas, J., Babcock, M.A., Gutmann, D.H., Korf, B., and Maria, B.L. (2009). Neurofibromatosis Type 1 Revisited. Pediatrics 123, 124–133.

Wong, V. (Wai C., and Lerner, E. (2015). Nitric oxide inhibition strategies. Futur. Sci. OA 1, fso.15.35.

Zerbi, V., Grandjean, J., Rudin, M., and Wenderoth, N. (2015). Mapping the mouse brain with rs-fMRI: An optimized pipeline for functional network identification. Neuroimage.

Zhu, Y., Romero, M.I., Ghosh, P., Ye, Z., Charnay, P., Rushing, E.J., Marth, J.D., and Parada, L.F. (2001). Ablation of NF1 function in neurons induces abnormal development of cerebral cortex and reactive gliosis in the brain. Genes Dev. 15, 859–876.

